## Supplementary information for "Structure of a Plant-Specific Partitivirus Capsid Reveals a Novel Coat Protein Architecture with a Hypervariable Protrusion"

Figure S1 – VLPs and SDS-PAGE

Figure S2 – Local resolution and FSC

Figure S3 – Monomer alignment and b-factors

Figure S4 – PCV coat protein organisation

Table S1 – Cryo-EM data collection, refinement and validation statistics

Figure S5 – Multiple sequence alignment of Deltapartitivirus coat proteins

Figure S6 – Alignment for all identified long-form CPs

Figure S7 – Disorder prediction for all identified long-form CPs

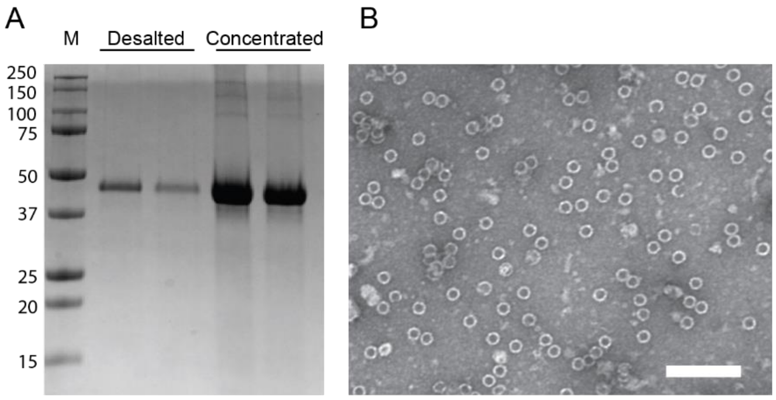


C

MGDRVNAQDDDTVVPHQAPLQPAALQQDLTRSADYLLDNVRIGNHRQRYDKYRRYVLLRS
SEIFTSLVAIYAHIFSSYWQHFRRFTDQFQAPTGVQLPTFVARVYISTWLHDLYCSIREA
TRSISPLAFNERYSYELLPYSTEYDPFLAFLSMSIKPTHIQHTPENTLWIPILCENYDWD
RNEANHNPFGITNFTLNSNLFYGLLAILKERKEFKLSTLTTNTIGRPCWLFDWHDNVQVC
AWFPREANFNSQDVTAAYIIGVACTPKLGPSDDDAWKYYASLNSVPTFTPTEPRLTNRRS
YGAYEVRTRETENNYFLPDSLLNIIEDFTATGTTQRRKIRRPSATSASTGAAIIIRDTPG
TASTATTSTTETEVTFPPVIRTKIRDWYYHSRVILELEDNSRTAALRMFIIA

**Figure S1.** Purification of PCV-1 VLPs. **(A)** SDS-PAGE of two gradient fractions from ultracentrifugation showing a single protein following desalting to remove iodixanol and following concentration using ultrafiltration. **(B)** Negative staining transmission electron microscopy of PCV-1 VLPs, bar = 200 nm. **(C)** Peptide coverage (74%) from mass spectroscopy analysis of bands cut from (A) shows presence of the N-terminal and internal disordered regions. Highlighted sequences (green) indicate peptide coverage and the underlined sequences show regions of disorder.

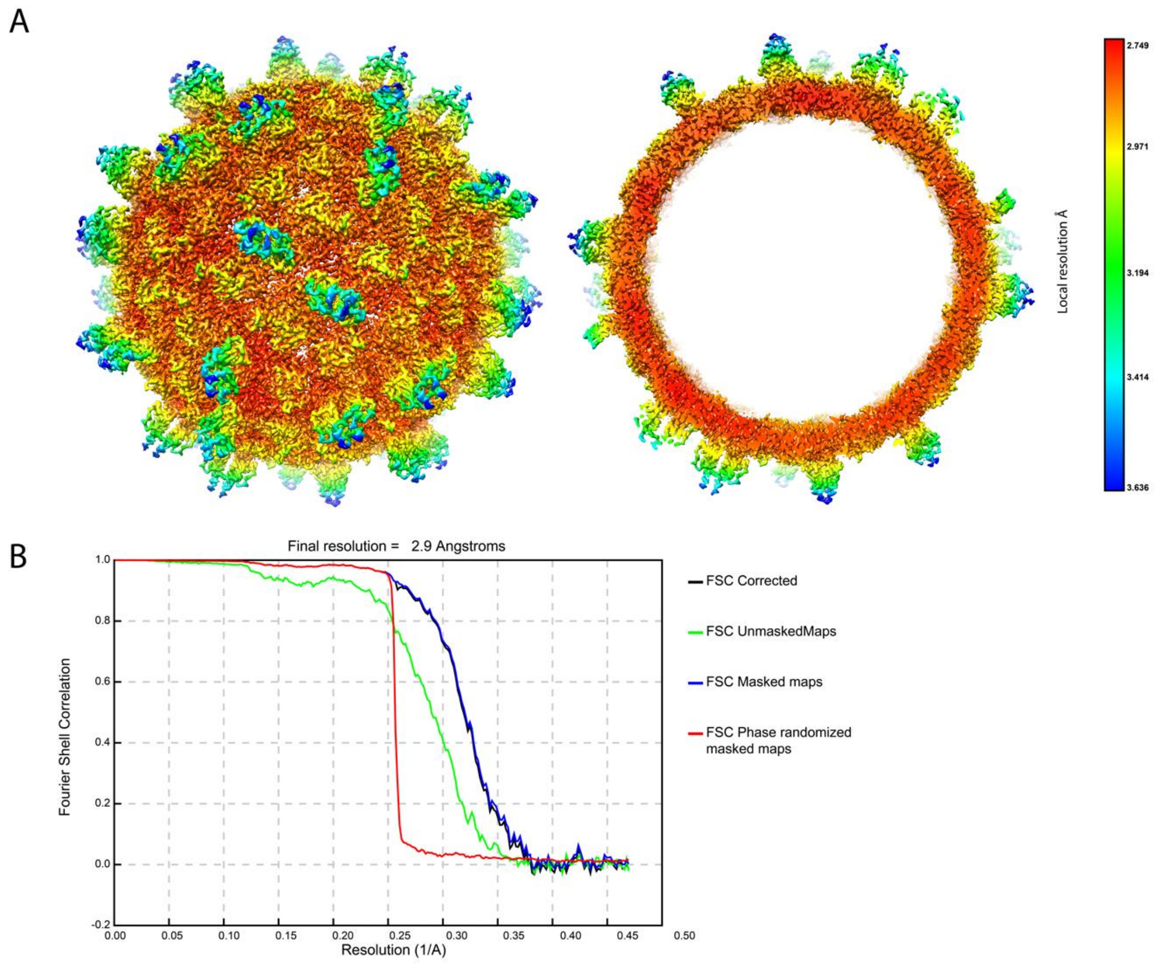


**Figure S2.** PCV-1 VLP reconstruction local resolution and Fourier shell correlation. **(A)** Local resolution filtered map of PCV-1 VLP coloured according to resolution, **(Left)** capsid surface, **(Right)** central slice. **(B)** Fourier shell correlation – the resolution that corresponds to an FSC coefficient of 0.143 is 2.9 Å.

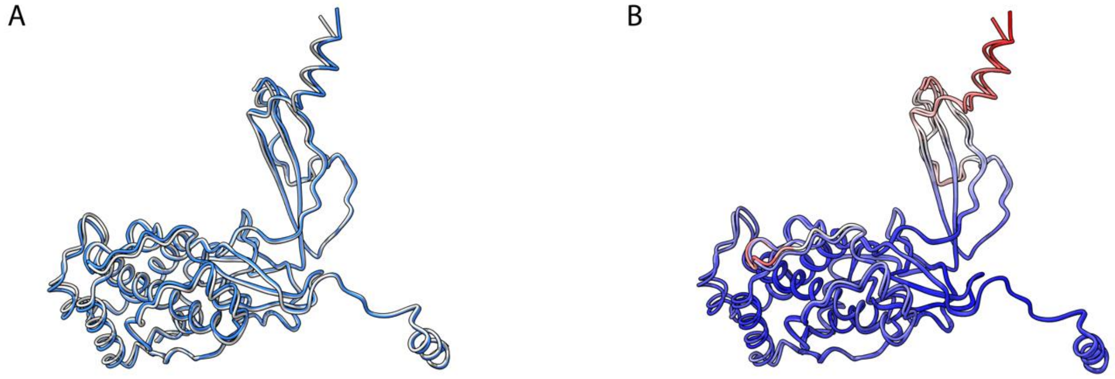


**Figure S3.** Superposition of chain A and chain B from a single asymmetric unit of PCV1. **(A)** Superposed PCV monomers, shown as backbone ribbons for clarity. Coloured according to chain with chain A coloured blue, and chain b coloured grey. **(B)** Superposed PCV monomers, shown as backbone ribbons for clarity. Coloured according to b-factor, from low (blue) to high (red).

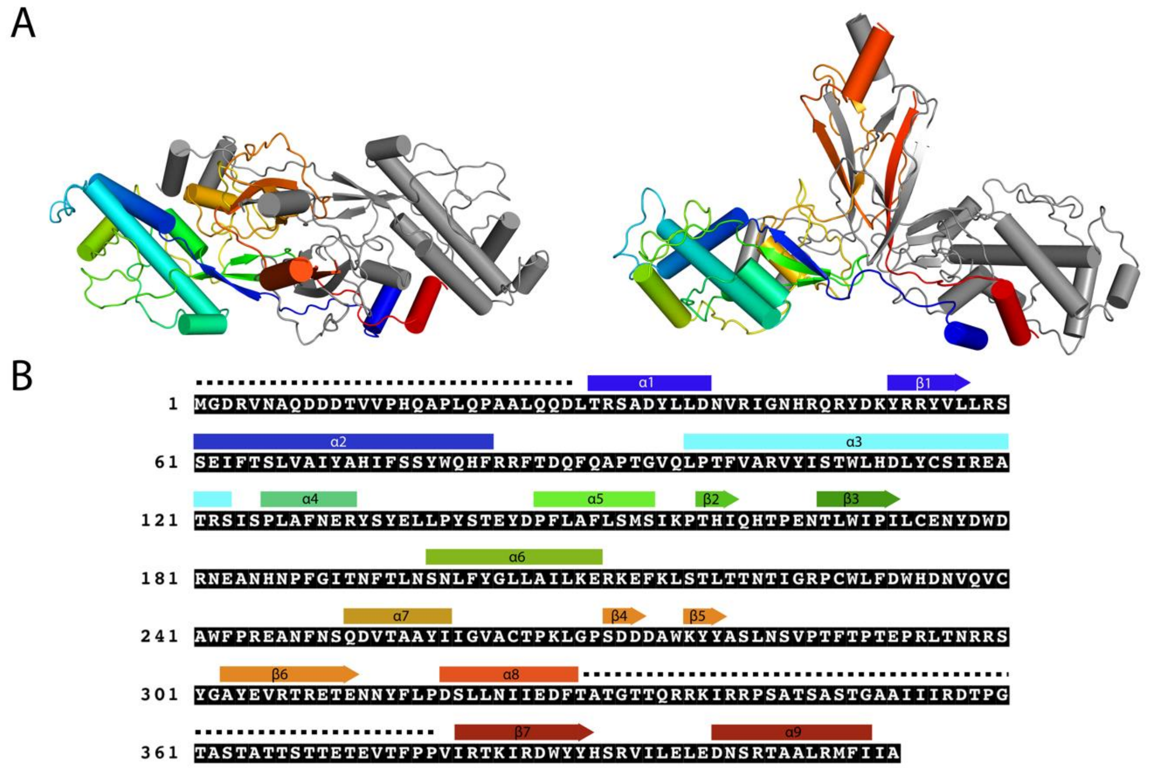


**Figure S4.** PCV coat protein composition. **(A)** PCV CP shown in cartoon form with monomer A coloured from N (blue) to C terminus (red). Monomer B is coloured grey for clarity. **(B)** PCV CP sequence labelled with secondary structure elements. Regions of disorder, that are not present in the final 3D reconstruction are denoted by a dashed line.

**Table S1.** Cryo-EM data collection, refinement and validation statistics

| | **PCV-1 VLP** |
|---|---|
| **Data collection and processing** | |
| Sample applications to grid | 1 |
| Magnification | 75, 000 x |
| Voltage (kV) | 300 |
| Electron exposure (e–/Å$^2$) | 74.26 |
| Defocus range of micrographs (μm) | -0.5 to -2.0 |
| Pixel size (Å) | 1.065 |
| Symmetry imposed | I |
| Initial particle images (no.) | 260, 000 |
| Final particle images (no.) | 105, 000 |
| Map resolution (Å) | 2.9 |
| FSC threshold | 0.143 |
| Number of frames | 59 |
| **Refinement** | |
| Map sharpening *B* factor (Å$^2$) | -127.7 |
| Model composition | |
| Protein residues | 668 |
| Nucleic acids | 0 |
| R.m.s. deviations | |
| Bond lengths (Å) | 0.006 |
| Bond angles (°) | 0.664 |
| Validation | |
| Clashscore | 8.29 |
| Poor rotamers (%) | 8.69 |
| Ramachandran plot | |
| Favored (%) | 91.36 |
| Allowed (%) | 8.18 |
| Disallowed (%) | 0.45 |

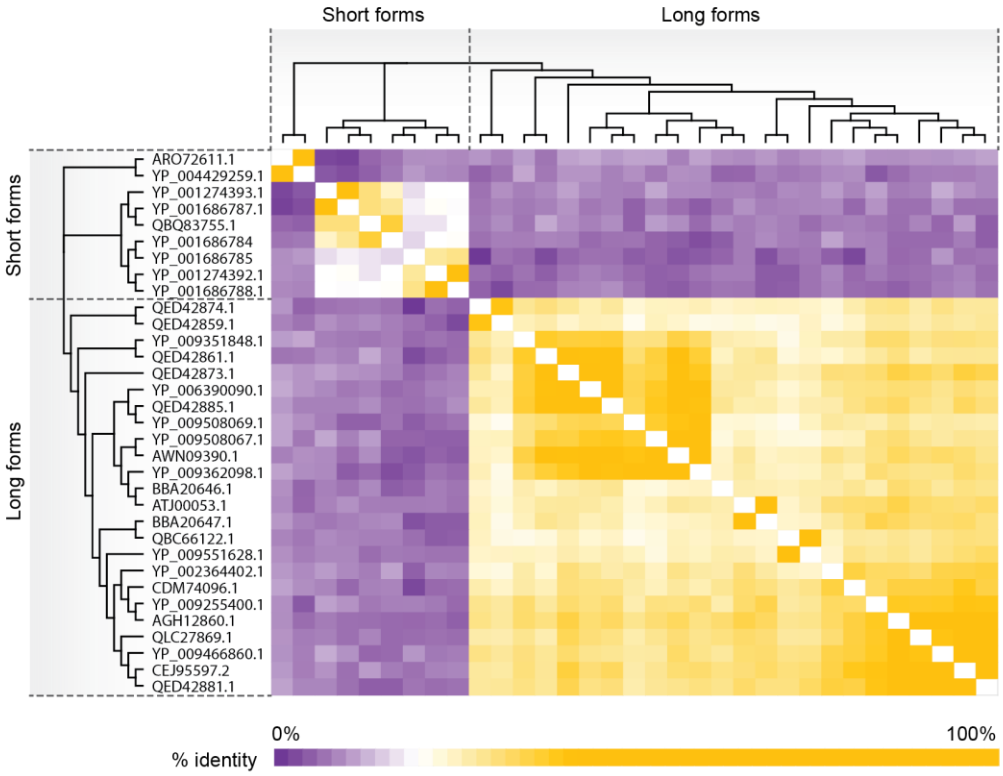


**Figure S5. Sequence comparison of Deltapartitivirus coat proteins.** Colour coded distance matrix based on a multiple sequence alignment performed using Clustal OMEGA (Sievers and Higgins, 2018) and the corresponding cladogram of the neighbour-joining tree. Sequence comparisons shows that there are two distinct forms of the coat protein among the deltapartitiviruses, a long form (374-483 aa) and a short form (335-348 aa). Between the long forms there is 20-91% pairwise identity with no more than 12% similarity with short any form coat proteins. Between the short form viruses there is 6-70% pairwise similarity.

There are five members of the deltapartitivirus (NCBI:txid1511810) recognised by the ICTV (Vainio et al., 2018). Three of these have the long form of the coat protein; Pepper cryptic virus 1 (PCV-1), Pepper cryptic virus 2 (PCV-2), and Beet cryptic virus 2 (BCV-2) that has two genomic segments encoding putative coat proteins. In addition, the ICTV notes four related viruses that are unclassified, of which one has the long form coat protein; Persimmon cryptic virus. The NCBI also lists three unclassified deltapartitiviruses (NCBI:txid1985162), of which one has the long form coat protein; Medicago sativa deltapartitivirus 1. PSI-BLAST using the PCV-1 coat protein identifies a further 18 viral sequences related to the long form coat proteins, and which are themselves long form coat proteins. The single exception to the length demarcation is a Citrullus lanatus partitivirus coat protein at 337 aa .

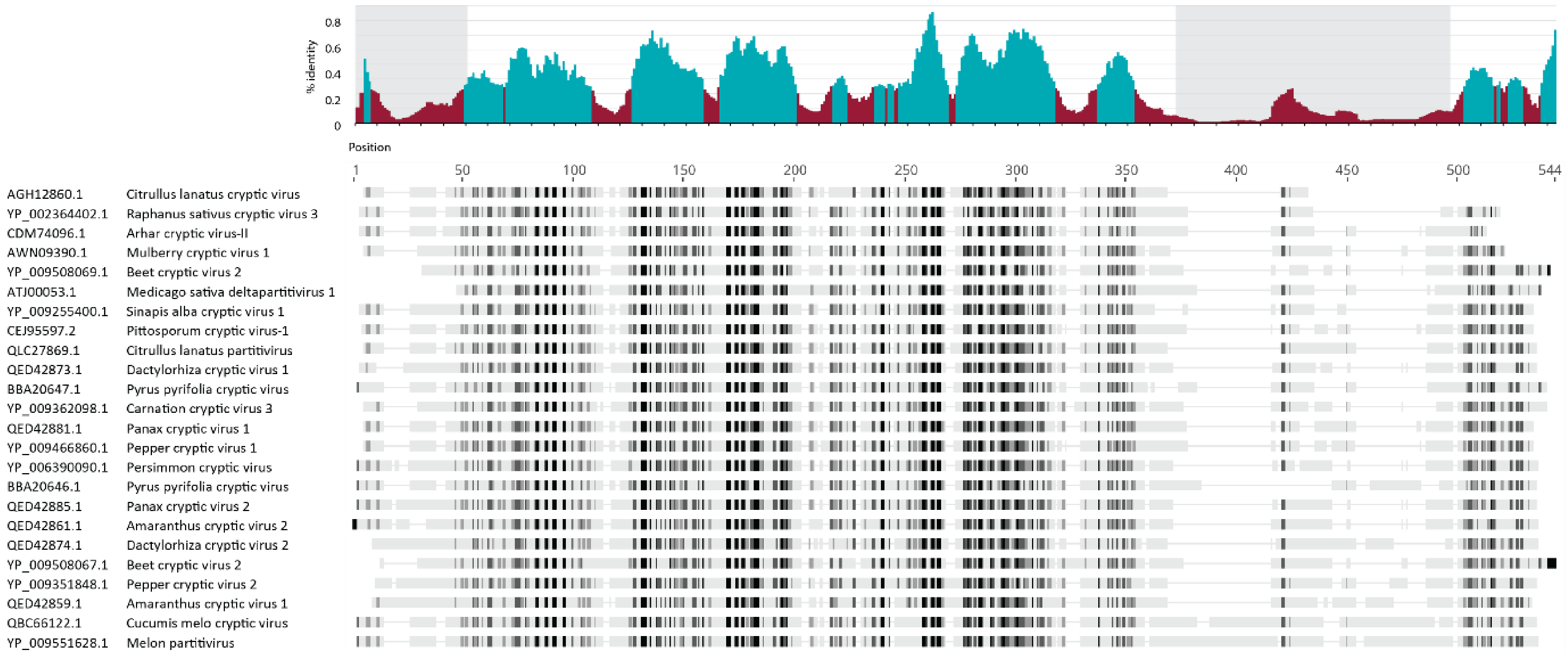


**Figure S6. Alignment of CP sequences of all 24 identified long form deltapartitivirus CP sequences.** Shaded regions correspond to the unresolved regions of PCV-1 CP and positions coloured cyan correspond to ≥ 25 % identity using a 10-position rolling average. The alignment is ordered according to the length of the CP sequence.

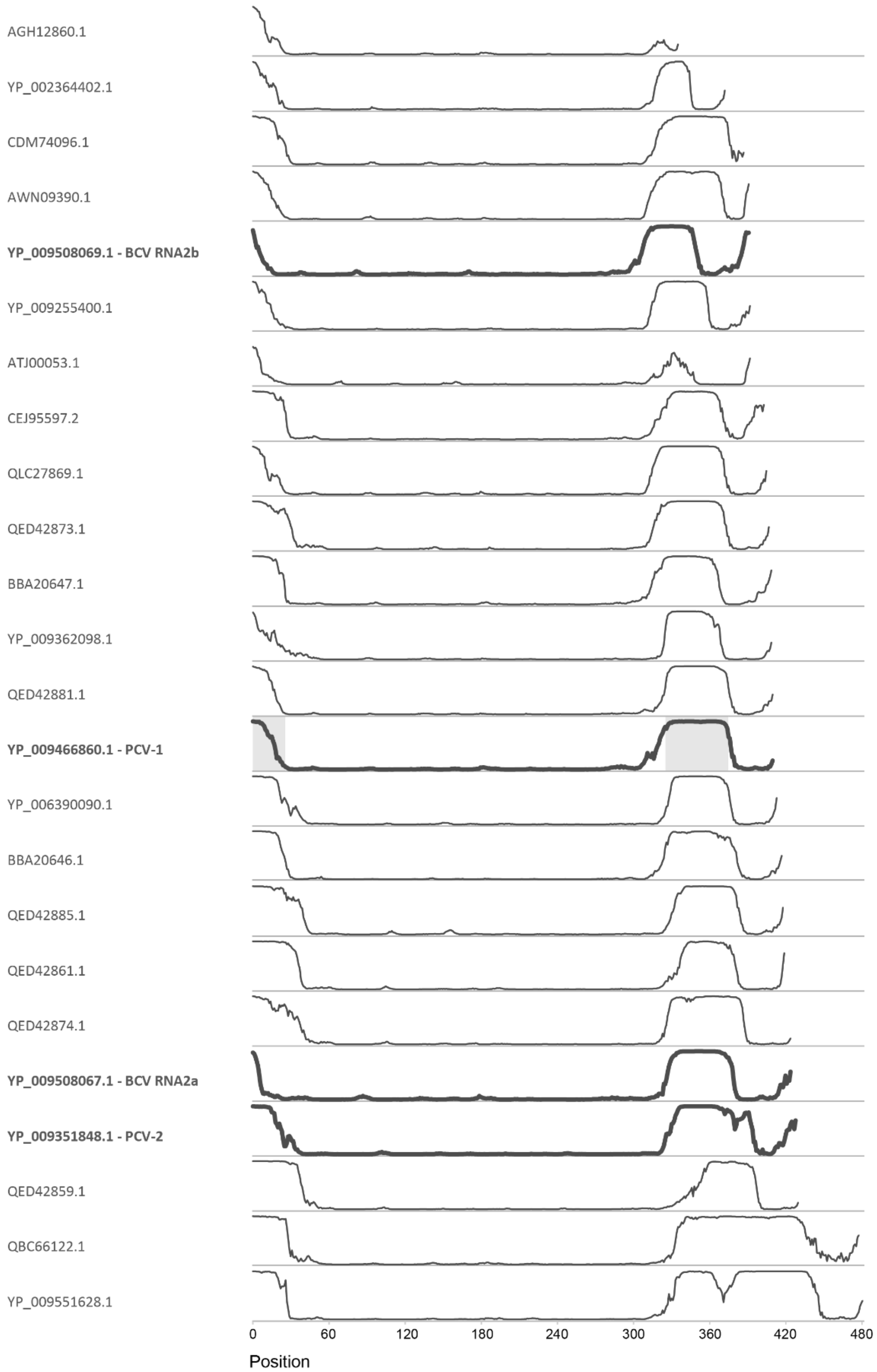


**Figure S7. Predicted disorder regions for the long form deltapartitiviruses.** Predictions using NetSurfP-2.0 (Klausen et al., 2019) show that the putative region of disorder on a scale 0 to 1 starts in approximately the same position for all 24 identified long form deltapartitivirus CP sequences, which are presented in order of the sequence length. Bold series represent those among the ICTV-recognised *deltapartivirus* genus (Vainio, Chiba et al. 2018) and the experimentally determined disorder for the PCV-1 CP is shaded.
